# Highly potent antisense oligonucleotides (ASOs) targeting the SARS-CoV-2 RNA genome

**DOI:** 10.1101/2022.11.28.518195

**Authors:** V. Dauksaite, A. Tas, F. Wachowius, A. Spruit, M.J. van Hemert, E.J. Snijder, E.P. van der Veer, A.J. van Zonneveld

## Abstract

Currently the world is dealing with the third outbreak of the human-infecting coronavirus with potential lethal outcome, cause by a member of the Nidovirus family, the SARS-CoV-2. The severe acute respiratory syndrome coronavirus (SARS-CoV-2) has caused the last worldwide pandemic. Successful development of vaccines highly contributed to reduce the severeness of the COVID-19 disease. To establish a control over the current and newly emerging coronaviruses of epidemic concern requires development of substances able to cure severely infected individuals and to prevent virus transmission. Here we present a therapeutic strategy targeting the SARS-CoV-2 RNA using antisense oligonucleotides (ASOs) and identify locked nucleic acid gapmers (LNA gapmers) potent to reduce by up to 96% the intracellular viral load *in vitro*. Our results strongly suggest promise of our preselected ASOs for further development as therapeutic or prophylactic anti-viral agents.

**One sentence summary:** ASOs (LNA gapmers) targeting the SARS-CoV-2 RNA genome have been effective in viral RNA (load) reduction *in vitro*.

## Main text

Recently the world was (and still in some areas is) dealing with the third outbreak of the human-infecting coronavirus with potential lethal outcome, caused by the member of the Nidovirus family, the severe acute respiratory syndrome coronavirus (SARS-CoV-2)^1^. Successful development of vaccines, especially the newly emerged mRNA-based vaccines, led to establish a control over the pandemic. However, antiviral drugs are and will be needed to complement vaccines in clinics when patients are already infected or as prophylactics for high-risk groups and to control the future outbreaks of coronaviruses. Multiple attempts to repurpose existing drugs resulted in few candidates to some extent still currently used in clinics (e.g., remdesivir, hydroxychloroquine). Multiple other tactics have been developed particularly aiming to reduce viral transmission, as e.g., the “miniprotein”^2^, lipopeptides^3^, antibody cocktails, etc. We took a parallel approach to develop antisense oligonucleotides that specifically target SARS-CoV-2 viral RNA, thereby blocking viral replication. Here we present selection and *in vitro* evaluation of new antivirals against SARS-CoV-2 with a potential to develop into a new medicine of innovative treatment and prevention options for emerging coronaviruses of pandemic concern.

The potential of small molecule drugs that target viral RNA components as antiviral agents has been widely recognized^4^. SARS-CoV-2 is positive-sense single-stranded RNA virus with a compact genome of approx. 30 kilobases (kb)^5^. Viral genome contains a 5’ untranslated region (5’UTR) with a 5’ leader sequence, almost two thirds of the genome occupying ORF1a/b region, encoding non-structural viral proteins and the downstream 3’ segment with sequences for the structural proteins (including spike (S), envelope (E), membrane (M) and nucleocapsid (N)) and a 3’ untranslated region (3’UTR)^5^. The features of the SARS-CoV-2 RNA genome are the non-coding 5’ and 3’ untranslated regions (UTRs) and the FSE site consisting of multiple highly conserved stem-loops/and more complex secondary structures (pseudoknots)^6–8^. Among the assigned functions to these structures is involvement of the 5’ UTR region in mediating viral replication and the FSE site, containing the programmed ribosomal frameshifting, pivotal for the ORF1ab translation^9^. Knowing the structural architecture of the SARS-CoV-2 RNA genome is essential in identifying (un)structured elements for the ASO designed and was available in multiple publications^10^.

We designed multiple viral RNA genome-targeting ASOs of variable chemical content (LNA gapmers, LNA-2’Ome gapmers, 2’Ome gapmers, 2’Ome’s, MOE’s, MOE gapmers). Gapmers are powerful tools for the mRNA loss-off-function studies as they catalyze RNase H-dependent degradation of complementary RNA targets, while the RNA ASOs are expected to interfere through the sterical hindrance. The LNA gapmers (containing the LNA flanking regions) are supposed to be superior to the other gapmers as the LNA region increases target binding affinity and confers nuclease resistance. All the ASOs contained fully modified phosphorothioate (PS) backbones, which ensure exceptional resistance to enzymatic degradation.

To facilitate ASO selection we conducted the initial functional evaluation in Vero E6 cell line using multiple SARS-CoV-2 genome sequences-containing Firefly luciferase reporter constructs without the use of infectious virus. For this assay, Vero E6 cells are transiently transfected with the virus genome-containing test construct, the control Renilla luciferase-containing construct and different ASOs. Luciferase readings are performed 30 h later. Virus genomic regions included in the reporters were: 5’ UTR (including the 5’ leader sequence) and the downstream ORF1a/b sequence (the beginning of the Nsp1); the FSE site of the ORF1a/b; coding sequences for the envelope (E), membrane (M) andnucleocapsid (N) proteins, the later together with the 3’ UTR sequence (depicted in Fig.1B). Each reporter construct tested resulted in selection of ASOs reducing the luciferase readings in comparison to no ASO/ctrlASO (scrambled sequence) controls (Fig.1C). At this time point, different ASO chemistries were tested and the LNA gapmers seemed to be more potent over the LNA-2’Ome or 2’Ome gapmers. One must note that this assay pre-selects the most potent ASOs for a particular reporter, but it is not possible to compare ASO potency between different reporters. As none of the tested RNA 2’Ome ASOs inflated high reduction activity, it raised a speculation that this mechanism is not testable in our assay set-up. To exclude potential effect resulting from ASO toxicity, cell viability assay was performed in parallel with this experiment (data not shown). Testing the most potent ASOs for the SARS-CoV-2 5’ UTR reporter in the same assay set-up in Calu-3 cell line (human lung cancer cell line) resulted in comparable reduction effects (Fig.S1).

**Figure 1.**
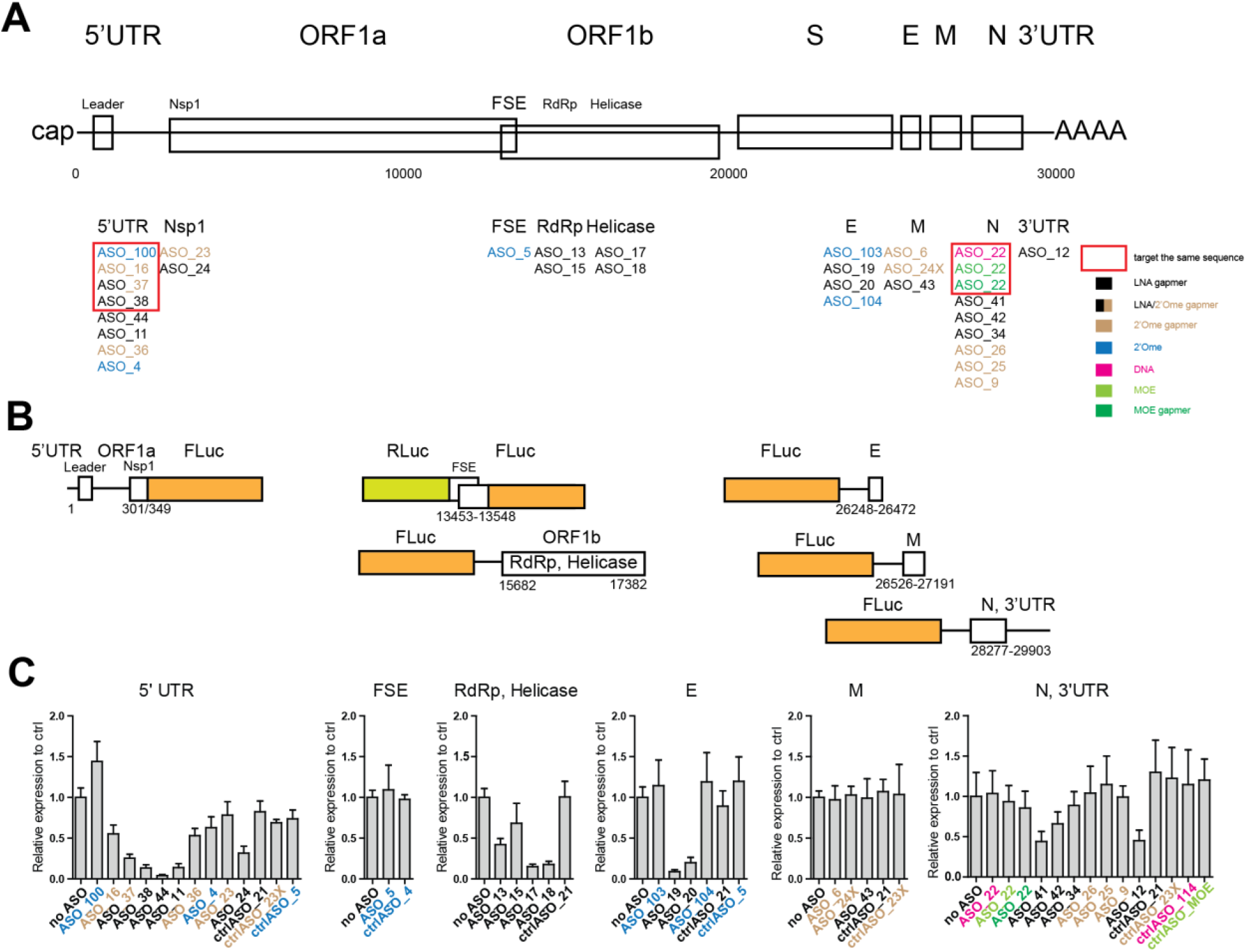
*In vitro* screening of ASOs targeting the SARS-CoV-2 genome in a cell-based assay. **A)** Structural overview of the SARS-CoV-2 RNA genome presented as a genic model showing the known organization of the genome (into the 5’ UTR, the two known ORFs (1a & 1b), major subgenomic RNAs and the 3’ UTR). Scale marker for the 30 kb genome is indicated. ASOs below are aligned according to the genomic regions they target. Legend to the ASO chemistry is indicated on the right. **B)** Schematic representation of luciferase (Firefly luciferase depicted in orange, Renilla luciferase depicted in yellow) reporter constructs used to screen ASOs in the *in vitro* assays. SARS-CoV-2 genomic regions, included in constructs, are indicated below. **C)** Vero E6 cells were transiently transfected with the reporter construct harboring the SARS-CoV-2 genomic regions, the control constructs and treated with the indicated ASO. Luciferase readings were assayed 30 h later. Inhibitory effect was calculated as the ration of relative luminescence units in the presence of a specific concentration of ASO and the relative luminescence units in the absence of ASO (no ASO) and corrected for background luminescence. Each ASO in a particular experiment was assayed in triplicate. Data are means (±) standard deviation (SD) from at least three independent experiments.

SARS-CoV-2 5’ UTR-targeting lead candidate ASOs were tested to select the most potent gapmer sequence. Different lengths of the LNA/DNA nucleotides in the gapmer were tested. Fig.2A and B show that the initial arrangement of nucleotides in the gapmer, central 10-13 DNA nucleotides flanked with the four LNA nucleotides from the 5’- and the 3’-ends, was the most optimal.

**Figure 2.**
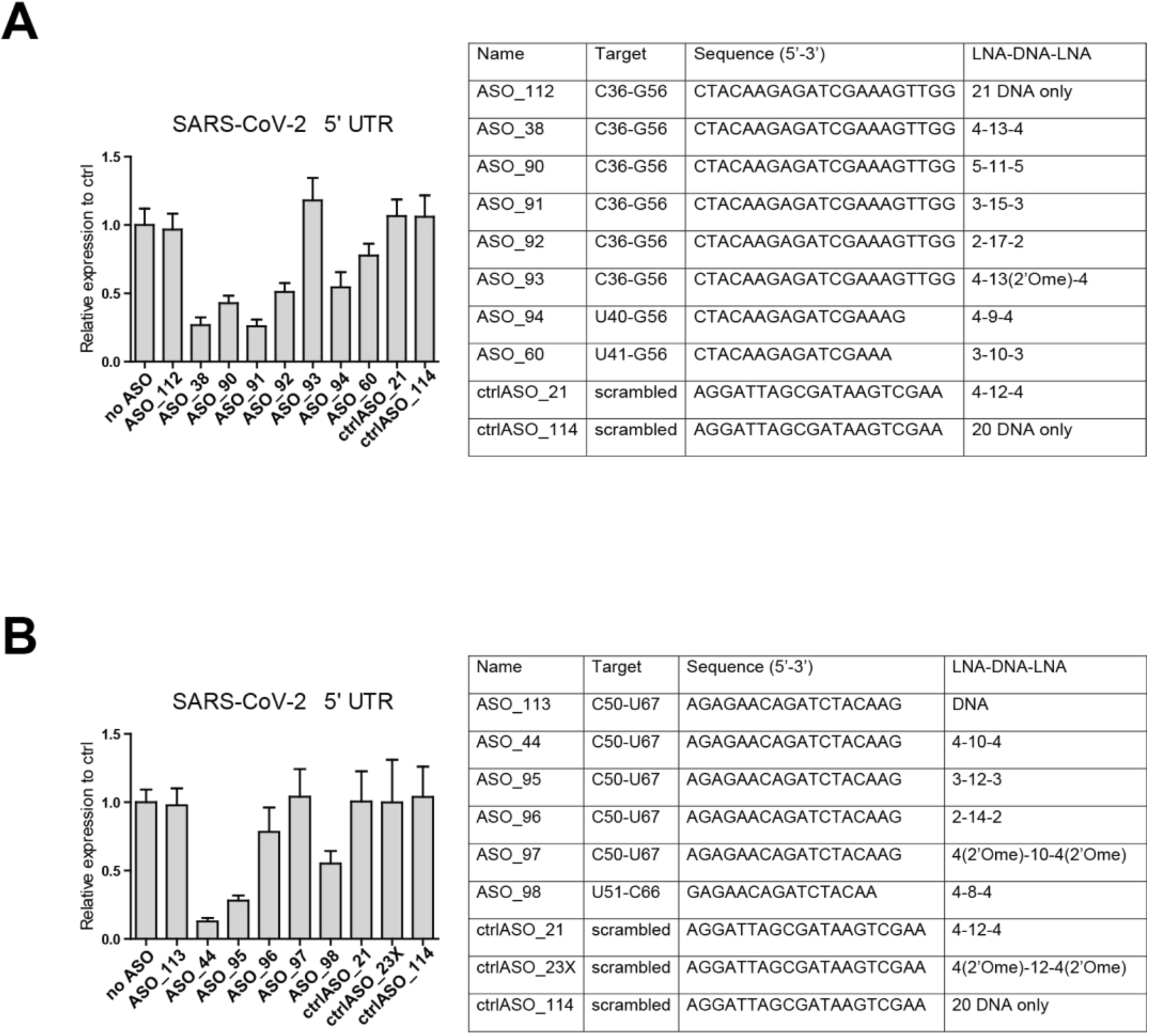
Lead candidate optimization. **A)** Vero E6 cells were transiently transfected with the reporter construct harboring the SARS-CoV-2 5’ UTR genomic region, the control constructs and treated with the indicated ASO. ASOs contain the same sequence (except the ones that are shorter) and differences introduced into the ASO chemistries are depicted in the table on the right. Luciferase readings were assayed 30 h later. Each ASO in a particular experiment was assayed in triplicate. Data are means standard deviation (SD) from at least three independent experiments. **B)** Vero E6 cells were transiently transfected with the reporter construct harboring the SARS-CoV-2 5’ UTR genomic region, the control constructs and treated with the indicated ASO. ASOs contain the same sequence (except ASO_98 that is shorter) and differences introduced into the ASO chemistries are depicted in the table on the right. Luciferase readings were assayed h later. Each ASO in a particular experiment was assayed in triplicate. Data are means standard deviation (SD) from at least three independent experiments.

To evaluate the actual viral repressive effect of ASOs selected in the reporter assays we performed experiments using live viral infection. Initially we used more commonly used experimental set-up when Vero E6 cells are first pre-treated (transfected) with ASOs for 8 h and then infected with live SARS-CoV-2 for 16 h (Fig.S2). However, after these initial experiments we switched to the more clinically relevant experimental set-up which would as well strengthen the potential translational use of identified ASO’s. In this case, Vero E6 cells are first infected for 1 h at 37°C with the live SARS-CoV-2 virus followed by virus removal and the 16-18 h ASO treatment of the infected cells. Intracellular RNA was assayed for the reduction in viral RNA loads in qRT-PCR using primers in viral RdRp and E genes. Using both approaches, the 5’ UTR-targeting ASOs 38 and 44 as well as the N gene-targeting ASOs 41 and 42 were identified as the most potent, leading to the 53-87% reduction in the intracellular viral load when used alone (depicted in Fig.3A). Using any combination of these ASO’s, their potency was retained, and the effect was even strengthened, leading up to 96% of the intracellular viral load reduction (Fig.3B). The MTT assay performed on these samples demonstrated no cell toxicity of the ASOs used (Fig.S3).

**Figure 3.**
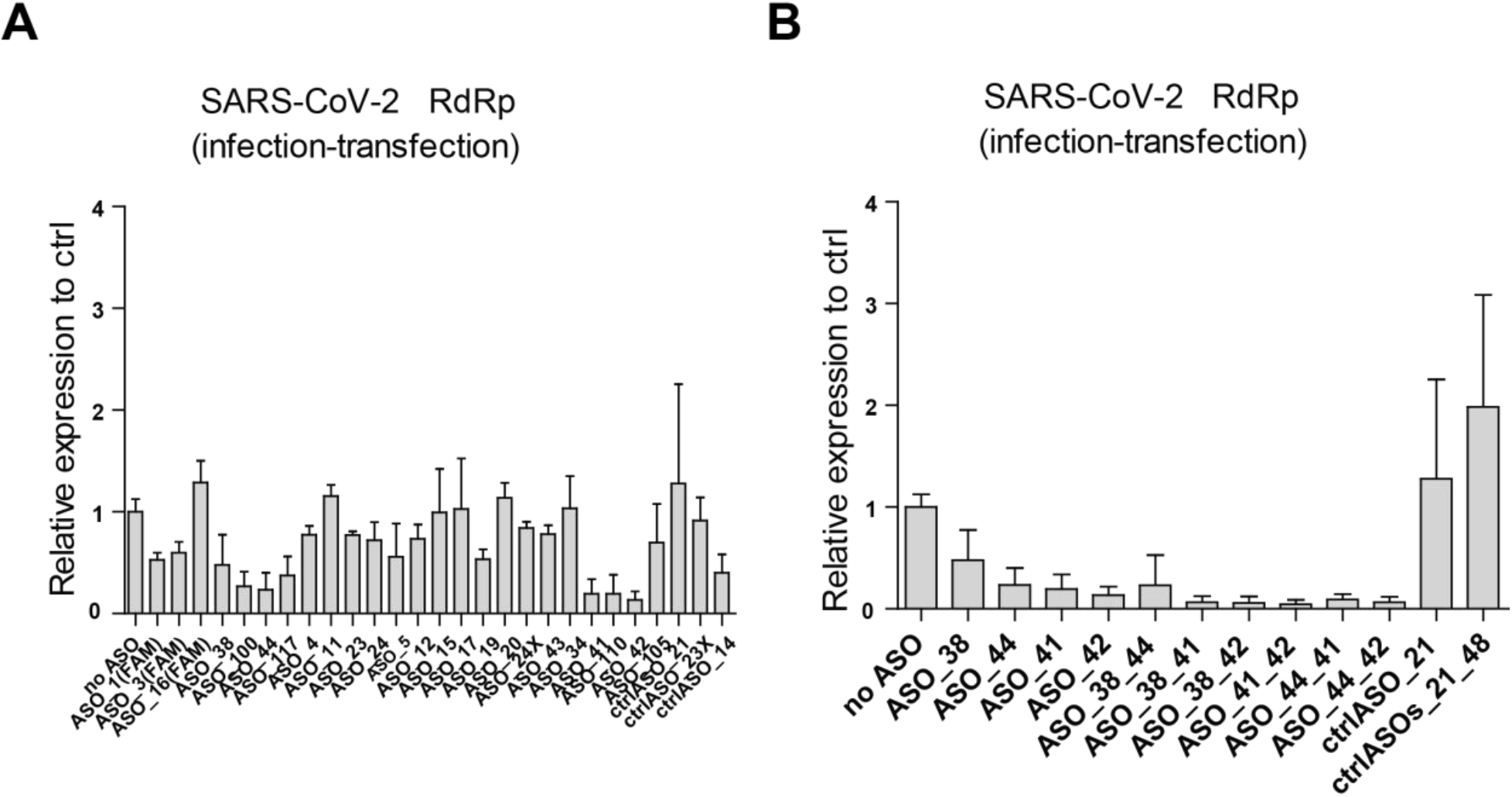
Selected ASOs are potent in reducing intracellular viral RNA levels. **A)** The most effective ASOs targeting each viral region have been further tested in transient transfections of the Vero E6 cells using the live SARS-CoV-2 virus (infection first, then ASO treatment (transfection)). Viral loads were detected via qRT-PCR. **B)** In the part A preselected lead candidates were further screened separately or in combination (synergy effect) in Vero E6 cells using the live SARS-CoV-2 virus and first infection then transfection approach. The percentage of intracellular viral load reduction is shown relative to the control (no ASO treatment). Data are means standard deviation (SD) from three independent experiments.

Additional reporter constructs have been generated to assay potency of lead candidate ASOs against highly conserved sequences from other betacoronavirus family members (Fig4B). To evaluate the potential for broad-spectrum activity, the lead candidate ASOs targeting the SARS-CoV-2 5’ UTR region were assayed on constructs carrying 5’ UTR sequences from SARS-CoV, MERS-CoV and MHV viruses. These constructs were created by fussing viral genome sequences starting from the first nucleotide to position in the ORF1a region (exact nucleotide positions indicated in Fig.4A) to the Firefly luciferase gene. Fig.4C shows the potential of lead candidate ASOs 38 and 44 to be most active against SARS-CoV viral sequences and in addition showed considerable potency against MERS and MHV sequences.

**Figure 4.**
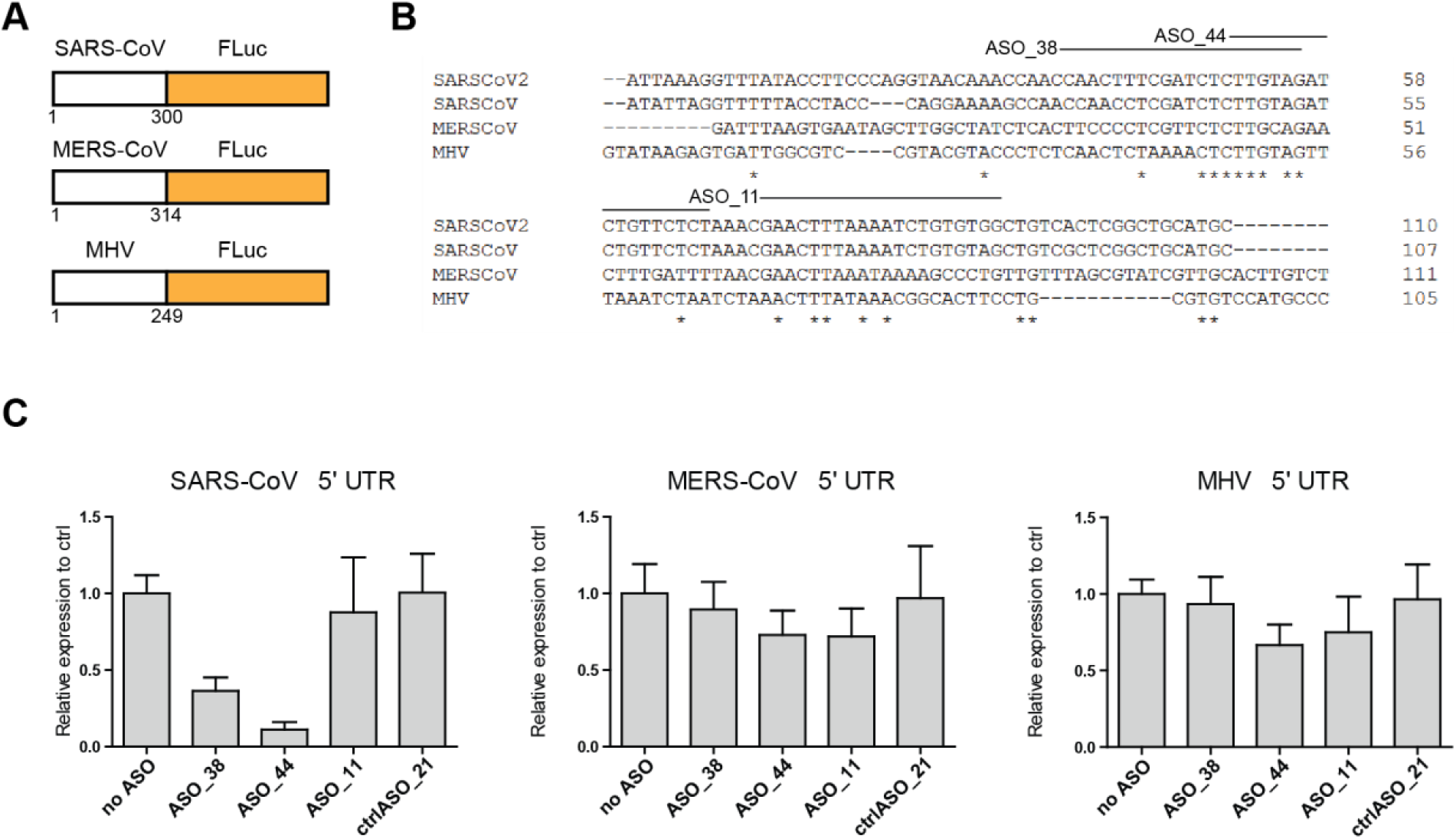
*In vitro* screening of SARS-CoV-2 genome-targeting ASOs against the other SARS-related betacoronaviruses. **A)** Schematic representation of the luciferase reporter constructs harboring the 5’UTR regions of SARS-CoV, MERS-CoV and MHV. **B)** Clustal-performed alignment of the conserved nucleotides are indicated with *. **C)** Vero E6 cells were transfected with the reporter constructs where SARS-CoV-2 genomic sequences are fussed to the Firefly luciferase gene and treated with ASO. Luciferase readings were assayed 30 h later.

Based on the *in vitro* results presented here we expect that selected ASOs will be effective in the *in vivo* experiments and possess viral load reduction. As well, our *in vitro* data suggests that ASOs might be effective against other emerging coronaviruses of the pandemic concern. In addition, testing of already identified highly conserved weakly structured regions within viral RNA genomes might aid the development of antiviral ASO therapeutics^10^. Work still needs to be done to define the best administration method (intranasal or systemic) and the uptake in the upper and lower respiratory tract, in the systemic circulation and organs. Selected ASOs need to be evaluated for the efficacy *in vivo*, stability and toxicity. Operating by a different mechanism, ASOs could be envisioned as additives to the existing/in development antivirals for intranasal prophylactic approach, preventing viral transmission.

## Author contributions

conceptualization/funding acquisition EvdV, AJvZ; experiments VD, AT, FW; writing – original draft VD; final version: all co-authors provided feedback to the final draft.

## Data and material availability

all data is available in the manuscript or the supplemental material.

## Materials and methods

### Cell culture

Vero E6 (African green monkey kidney) cells were grown in Dulbecco’s modified Eagle’s medium (DMEM with GlutaMAX™, Gibco™) supplemented with 10% fetal bovine serum (FBS) in 5% CO2 at 37°C.

### Molecular cloning

To generate pCI-neo-FLuc reporter, FLuc gene was PCR-amplified and cloned into the pCI-neo plasmid which subsequently was used as a parental plasmid to generate SARS-CoV-2 sequences-containing reporter constructs.

### ASO design

All ASOs were produced by Eurogentec. ASO sequences and target locations in SARS-CoV-2 are specified in Supplementary TableS1 and represented schematically in Fig.1A.

### Transient transfections

Vero E6 cells have been seeded at a defined density a day before on the 24-well plate. Next day medium was removed and substituted with a fresh medium. Lipofectamine 3000 (Invitrogen) transfection mixes contained the reporter plasmid, the control plasmid expressing Renilla luciferase and ASO at the indicated concentration. After 6 h transfection mixes were removed and substituted with a fresh medium. After 30 h cells were lysed and luciferase readings performed using the Promega kit according to the manufactures instructions. Readings were performed using SpectraMax i3x device (Molecular Devices). Inhibitory effect was calculated as the ration of relative luminescence units in the presence of a specific concentration of ASO and the relative luminescence units in the absence of ASO (no ASO) and corrected for background luminescence. Each ASO in a particular experiment was assayed in triplicate. Data are means (+/-) standard deviation (SD) from at least three independent experiments.

### Transfections using live virus

Vero E6 cells have been infected for 1 h at 37°C with 12000 PFU/well of the live virus followed by the 16 h ASO treatment. After approx. 18 h post infection cell lysates were collected, and RNA isolation was performed using the standard RNA isolation protocol from Medical Microbiology. Multiplex qRT-PCR analysis was performed using Taqman probes against SARS-CoV-2 RdRp gene, PGK1 housekeeping gene was used for normalization.

### Cell toxicity assay

Vero E6 cells have been infected (see above) followed by the 16 h ASO treatment. After approx. 18 h post infection, supernatants were collected and used to perporm a cell proliferation assay (MTS) according manufactures (Promega) instructions.

## Supplementary Materials

List of Supplementary Materials: Figures S1–S3 and supplemental TableS1.

## Supplementary figures

**Figure S1.**
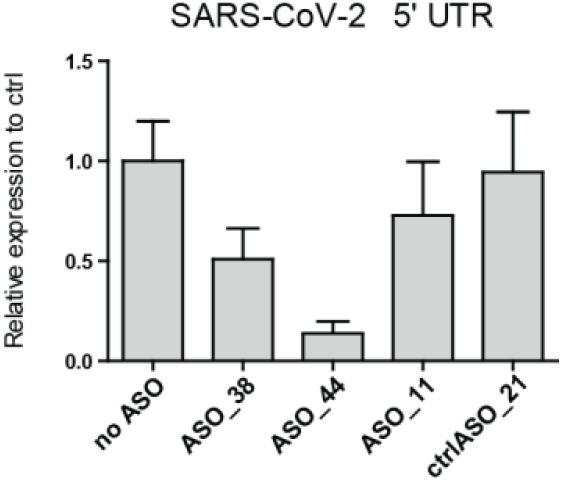
Selected ASOs targeting the SARS-CoV-2 5’ UTR region demonstrate comparable reduction in luciferase expression in a lung cell line. Calu-3 cells were transiently transfected with the reporter construct harboring the SARS-CoV-2-5’ UTR, the control construct and treated with the indicated ASO. Luciferase readings were assayed 48 h later. Each ASO in a particular experiment was assayed in triplicate. Data are means standard deviation (SD) from at least three independent experiments.

**Figure S2.**
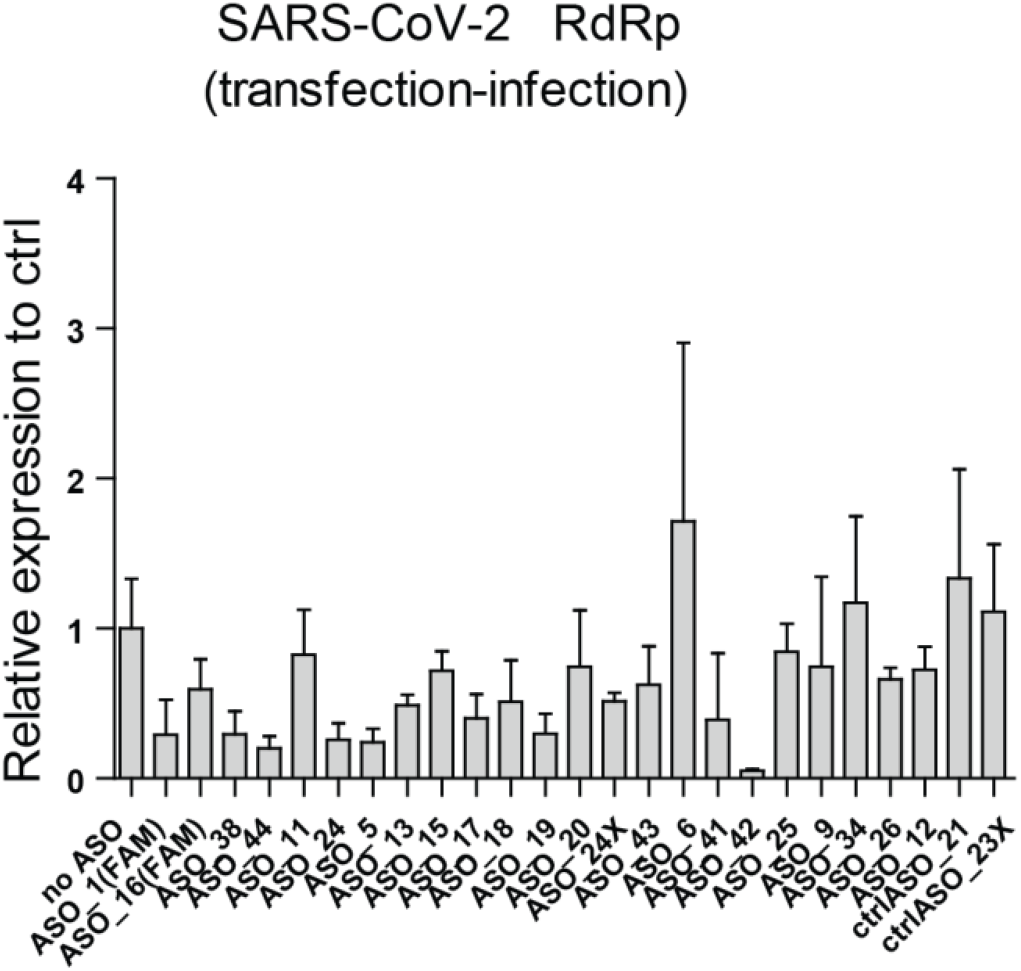
Selected ASOs are potent in reducing intracellular viral RNA levels in transfection first, infection second experimental set-up. The most effective ASOs targeting each viral region have been further tested in transient transfections of the Vero E6 cells using the live SARS-CoV-2 virus (ASO treatment (transfection) first, infection second). Viral loads were detected via qRT-PCR. The percentage of intracellular viral load reduction is shown relative to the control (no ASO treatment). Data are means standard deviation (SD) from three independent experiments.

**Figure S3.**
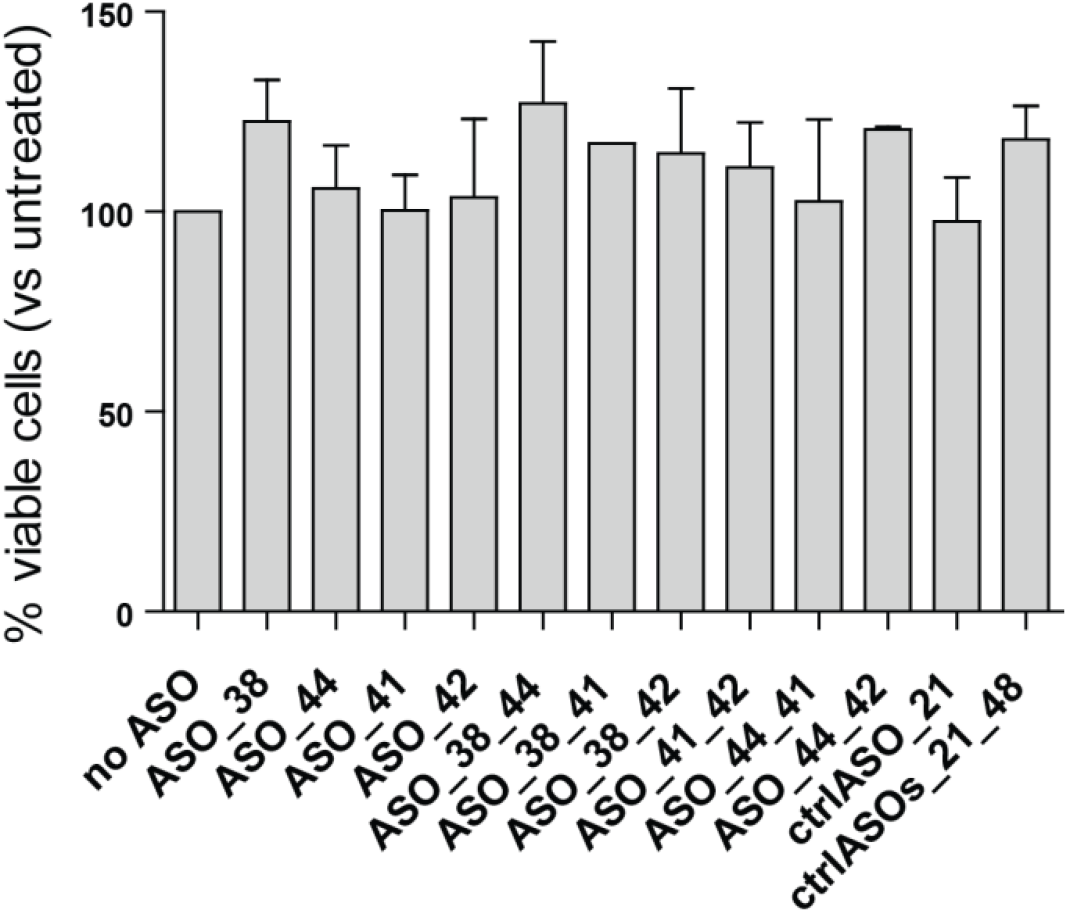
MTT assay summary for infection-transfection (synergy effect) experiments.

## Supplemental Table 1

**Table S1:**
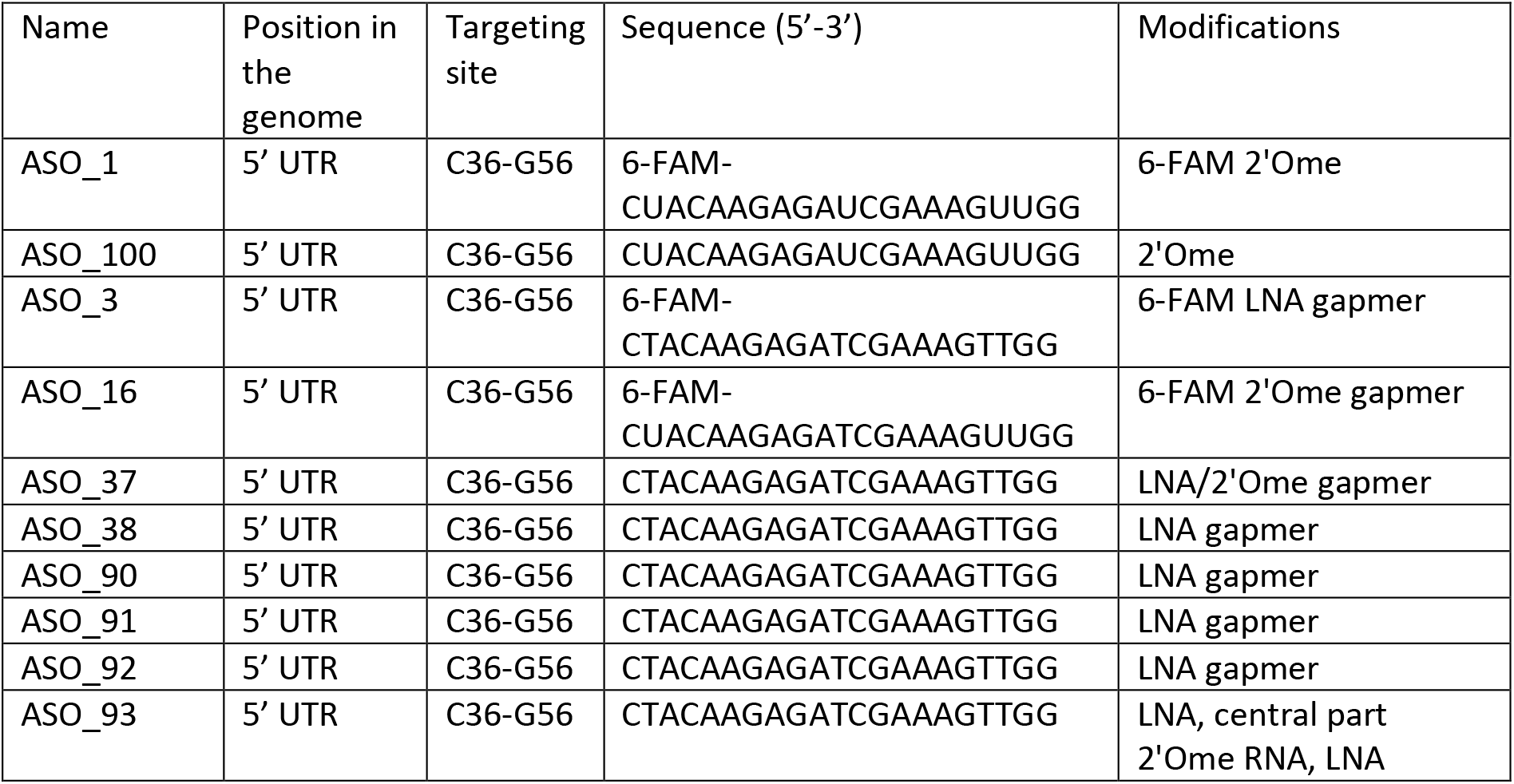
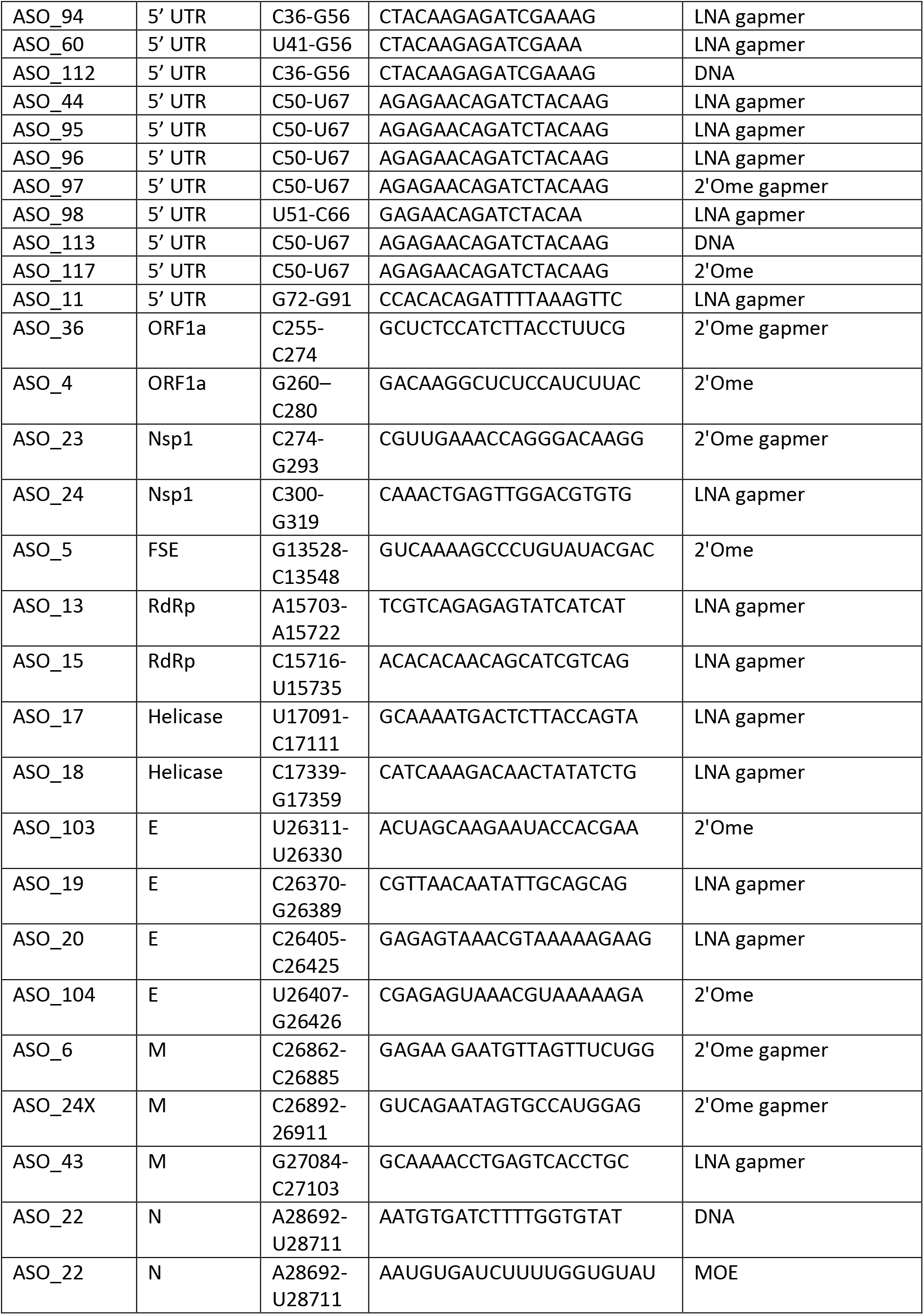
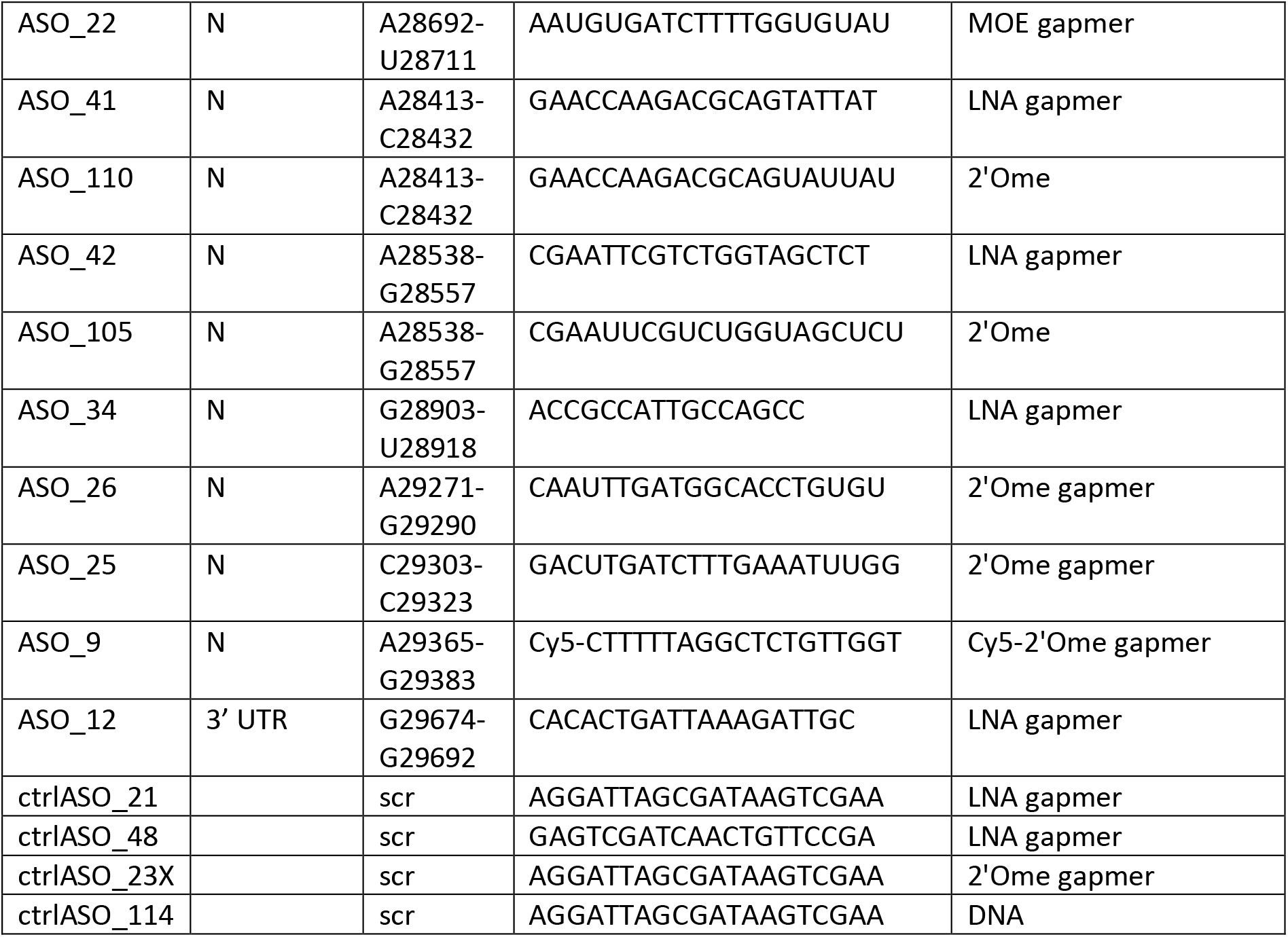
List of ASOs targeting the SARS-CoV-2 genome

